# Post-translational modifications distinguish amyloid-β isoform patterns extracted from vascular deposits and parenchymal plaques

**DOI:** 10.1101/2025.06.11.658821

**Authors:** Srinivas Koutarapu, Kaleigh F. Roberts, Reid A. Coyle, Jogender Mehla, Chihiro Sato, Gregory J. Zipfel, Randall J. Bateman, Katherine E. Schwetye, Soumya Mukherjee

**Affiliations:** Department of Pathology & Immunology, Division of Neuropathology, Washington University School of Medicine, St. Louis, MO, 63110, USA; Tracy Family SILQ Center, Washington University School of Medicine, St. Louis, MO, 63110, USA; Department of Neurology, Washington University School of Medicine, St. Louis, MO, 63110, USA; Taylor Family Department of Neurosurgery, Washington University School of Medicine, St. Louis, MO, 63110, USA

**Keywords:** Amyloid β (Aβ), Alzheimer’s disease (AD), cerebral amyloid angiopathy (CAA), parenchymal plaques, mass spectrometry

## Abstract

Deposition of amyloid-β (Aβ) aggregates is a core pathological hallmark of both cerebral amyloid angiopathy (CAA) and extracellular parenchymal plaques in Alzheimer’s disease (AD). While both disease processes share progressive, decades-long deposition of fibrillar Aβ peptide, they differ in isoform composition. We hypothesized that post-translational modifications (PTMs) on Aβ would also differ between CAA and parenchymal plaques. Using Lys-N enzymatic digestion followed by quantitative mass spectrometry, we profiled Aβ isoforms and N-terminus PTMs (aspartic acid isomerization and pyroglutamate formation) across CAA severity and compared them to parenchymal plaque Aβ in AD. Moderate to severe CAA primarily featured intact N-terminus (Aβ_1-x_) (∼95%) with minimal N-truncated species (Aβ_2-x_, Aβ_3pGlu-x_, Aβ_4-x_), whereas parenchymal plaques displayed diverse N-terminus truncations and PTMs. Increasing CAA severity correlated with a shift from longer, hydrophobic C-terminal isoforms (Aβ_41_, Aβ_42_, Aβ_43_) to shorter, less hydrophobic C-terminal isoforms (Aβ_37_, Aβ_38_, Aβ_39_, Aβ_40_). Importantly, moderate and severe CAA displayed minimal isomerization of Asp-1 and Asp-7 residues, which correlated significantly (r > 0.9) with shorter C-terminal isoforms (Aβ_37_, Aβ_38_, Aβ_39_, Aβ_40_). These patterns suggest distinct Aβ aggregation mechanisms in CAA versus parenchymal plaques. We propose that the intact N-terminus found in CAA with limited Asp isomerization is due to its inclusion within the protofibril structure (less disordered and inaccessible to PTMs), unlike the parenchymal plaques, where the N-terminus is more disordered and accessible to PTMs. These biochemical differences may reflect distinct protofibril architectures with potential implications for biomarker development for early CAA detection and therapeutic targeting of vascular and parenchymal Aβ.

## Introduction

Pathological hallmarks of Alzheimer’s disease (AD) are extracellular plaques and neurofibrillary tangles (NFTs)(1, 2); cerebral amyloid angiopathy (CAA)(3–5) is also observed in 85-95% of AD patients. Fibrillar amyloid-β (Aβ) is the main constituent of extracellular amyloid plaques of AD and vascular deposits of CAA(1, 6). Initial Aβ deposition in CAA starts within the basement membrane and then the tunica media of cerebral and leptomeningeal vessels. Extensive Aβ deposition in the vasculature may lead to fibrinoid necrosis and microaneurysm formation in the most severe cases(7). This ultimately compromises vascular integrity and increases the risk of micro- and lobar hemorrhages. CAA has been linked to increased amyloid-related imaging abnormalities (ARIA) in patients receiving anti-amyloid targeting therapies (20-40%, depending on the drug and clinical trial design)(8, 9). Understanding the underlying molecular signatures of CAA is crucial for developing better therapeutic strategies for targeting CAA as well as the development of specific biomarkers to predict which AD patients receiving anti-amyloid therapies are at higher risk of adverse side effects.

Biochemically, Aβ peptides that deposit as aggregates in the brain are usually 38−43 amino acids long and result from the sequential cleavages of the amyloid precursor protein (APP) by β-secretase and γ-secretase enzymes(10). N-terminal and C-terminal processing of Aβ results in the formation of various proteoforms with altered physicochemical properties(11), often changing seeding and aggregation propensity(12) and contributing to the formation of pathological deposits(13). While Aβ_1-40_ and Aβ_1-42_ are the most common isoforms found in CAA and parenchymal plaques of AD(14–16), respectively, many other Aβ isoforms have been identified in postmortem AD patients(17, 18). Additionally, multiple post-translational modifications (PTMs) of Aβ have been characterized, which further diversify the proteoforms of Aβ contributing to these pathologies(19, 20). The majority of Aβ PTMs extracted from the plaques are localized on the N terminus, including spontaneous non-enzymatic isomerization of the aspartic acid residues (Asp1, Asp7 and Asp 23)(6), pyroglutamate formation(20–22), and arginine citrullination(23, 24).

Moreover, Aβ mutation at position Asp7 (D7N) ― referred to as Tottori mutation and Aβ mutation at Asp23 (D23N) ― referred to as Iowa mutation, enhances the rate of oligomerization resulting in faster aggregation kinetics(25, 26). Interestingly, isomerization at Asp7 is abundant in both parenchymal plaque and amyloid deposited vasculature(14, 27, 28), while isomerization at Asp23 is mainly associated with heavy vascular amyloidosis associated with dementia and intracerebral hemorrhages(29). Substantial isomerization at Asp1 and Asp7 has been characterized from parenchymal plaque-derived Aβ(24, 30), (> 90% in AD and ∼ 50% in preclinical AD)(31). The degree of Asp1 and Asp7 isomerization is related to the age of the aggregates, suggesting that the parenchymal plaque Aβ accumulate these modifications over time(6, 19, 32). However, to what degree these Asp residues are modified in CAA remains currently unexplored. As these PTMs have been shown to alter physiochemical properties, including toxicity(32, 33), characterization of these changes and how they differ between CAA and parenchymal plaques in AD is critical for understanding the pathogenesis of these diseases to better predict response to anti-amyloid therapeutics and reduce the risk for adverse side effects such as ARIA.

In this study, we utilized a clinically well-characterized frozen human brain tissue cohort to investigate Aβ proteoform distribution across neuropathologic examination. Tissue was scored by an expert neuropathologist (none, mild CAA, moderate CAA and severe CAA) for both leptomeningeal and parenchymal vessels, accounting for density, circumferential accumulation, and dyshoric features. We isolated cortical vessels using density gradient centrifugation and separated the vascular proteins into soluble and insoluble fractions. We used immunoprecipitation coupled with liquid chromatography mass spectrometry (IP/MS) to analyze the Aβ isoform distribution in soluble and insoluble cerebral vessel fractions. We determined the heterogeneity of the N- and C-terminal isoforms across the severities of CAA and compared them to Aβ derived from insoluble parenchymal plaques from AD. In addition to a distinct Aβ isoform distribution in vascular versus parenchymal plaques, we also characterized the post-translational modifications (PTMs) including aspartic acid isomerization (Asp1 and Asp7), N-terminal pyroglutamate formation (pGlu3) and arginine citrullination (Arg5). The results indicate that origin of the N-and C-terminal heterogeneity and PTMs in vascular versus parenchymal plaque Aβ could be a result of the protofibril structure of the Aβ aggregates depositing in these two brain compartments. Our results have implications for better therapeutic strategies for targeting CAA as well as diagnostic biomarker development.

## Results

To investigate the differences in the biochemical composition of Aβ across severities of CAA conditions and parenchymal plaques, we first estimated the concentration of various Aβ species as well as their relative abundance to the total Aβ extractable from these brain compartments. To achieve this, we employed targeted mass spectrometry (MS/MS) analysis on insoluble and soluble fractions of CAA-laden purified vascular extracts to probe into the composition of Aβ with a specific focus on N-terminal and C-terminal species in mild, moderate, and severe CAA. We compared the proteoform distributions to controls without CAA and to parenchymal Aβ plaques. In most analyses, the insoluble and soluble fractions (Figure S2) showed similar proteoform distributions.

### Distinct N-terminal and C-terminal isoforms in CAA and parenchymal plaques

To compare the differences in the Aβ N-terminal truncations, we first analyzed the insoluble fraction of CAA-enriched vessels. We found that Aβ_1-15_ is the primary N-terminus in the insoluble fraction (Figure 2. Aa). Both the moderate and severe CAA groups showed significantly higher absolute concentrations of Aβ_1-15_ than the group without CAA, while the mild CAA group was not significantly different from controls. Quantitation of additional N-terminal isoforms (_Aβ2-15_, Aβ_4-15_, Aβ_3pGlu-15_, cit-Aβ_3pGlu-15_) showed concentrations over 100 times lower than Aβ_1-15_ in CAA-positive vessels. These additional isoforms followed a similar pattern wherein only moderate and severe CAA showed significantly higher levels than the control CAA-none group except for cit-Aβ_3pGlu-15_ (Figure 2. Ab, 2. Ac and 2. Ae).

**Figure 1.**
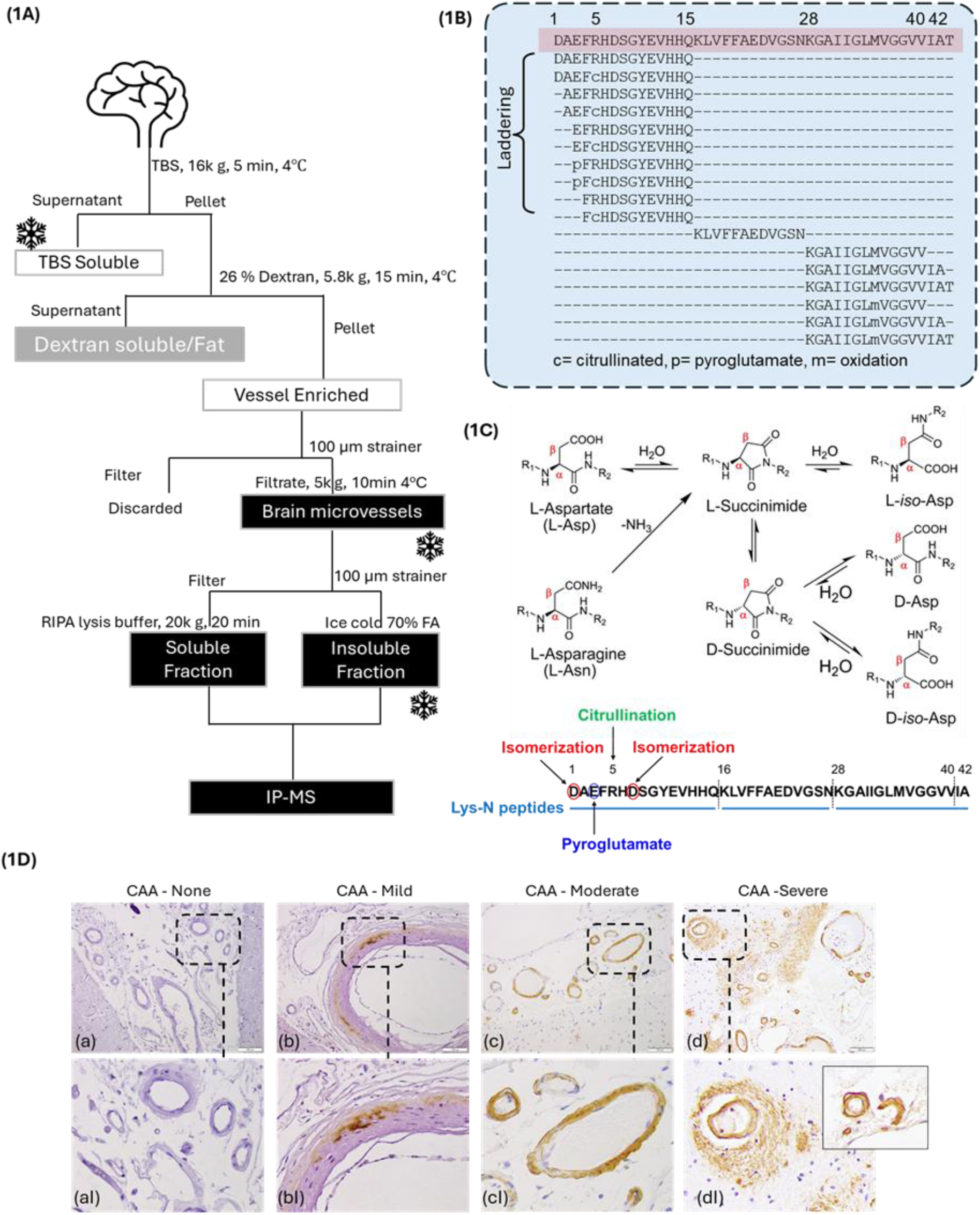
(1A) Schematic representation of the methodology for the extraction of Aβ from frozen occipital cortex tissue prior to IP/MS. (1B) Schematic diagram of Aβ peptide sequence showing various N- and C-terminal isoforms detected in AD brains using mass spectrometry (1C) Schematic representation of the mechanism of dehydration of L-Asp (native form) and deamidation of L-Asp resulting in a succinimide intermediate that subsequently leads to the isomerization/racemization after ring opening to D/L-iso-Asp and D-Asp. Schematic diagram of the Aβ peptide sequence showing sites of common PTMs investigated in this report. (1D) Representative immunohistochemistry of Aβ antibody (10D5) stained brain sections from neuropathologically assessed cases annotated as (1Da) CAA-none, (1Db) CAA-mild, (1Dc) CAA-moderate, and (1Dd) CAA-severe. (1DaI-dI) Magnified pictograms that depict the presence of sparse, patchy, and dyshoric CAA, in mild, moderate and severe CAA, respectively. Insert in (dI) represents the double-barreled CAA morphology representing severe CAA.

**Figure 2.**
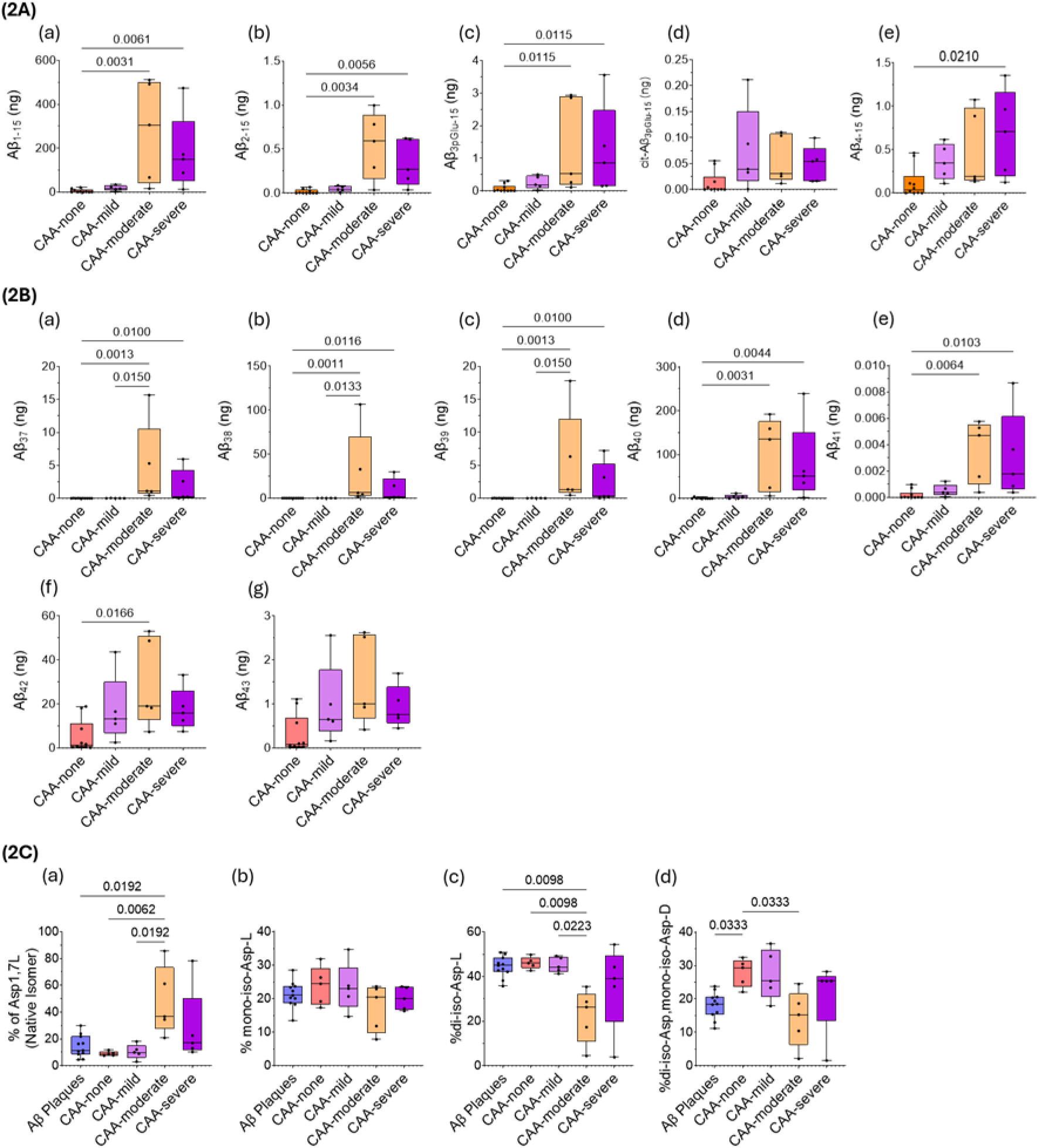
Isoform distribution of N-terminal Aβ proteoforms and C-terminal Aβ proteoforms. Box plots represent the Aβ isoform distribution of (2A) N-terminal Aβ proteoforms, (2B) C-terminal Aβ proteoforms, and (2C) Aβ1-15 isomers (Asp-1, Asp-7) in parenchymal plaques and CAA. Specifically, of the (2A) N-terminal Aβ proteoforms obtained from the insoluble fraction (2Aa) Aβ_1-15_, (2Ab) Aβ_2-15_, (2Ac) Aβ_3pGlu-15_, (2Ad) cit-Aβ_3pGlu-15_, and (2Ae) Aβ_4-15_, Aβ_1-15_ is the dominant species in CAA-moderate and CAA-severe cases. (2B) Among C-terminal forms Aβ_37_, Aβ_38_, Aβ_39_, Aβ_40_, Aβ_41_, Aβ_42_, and Aβ_43_ (2Ba-g), Aβ_40_ peptide is the dominant species, along with significant contribution from other C-terminal species in the order of Aβ_42_, Aβ_38_, Aβ_37_, and Aβ_39_. (2C) Differences in the distribution of Aβ_1-15_ isomers (Asp-1, Asp-7) between parenchymal plaques and CAA. P values were adjusted for multiple comparison false discovery rate (p<0.05).

Next, we investigated the truncations on the C-terminus Aβ in the vascular insoluble fraction. We found that the concentrations of Aβ_37_, Aβ_38_, Aβ_39_, and Aβ_40_ in the brain microvessels increase with the severity of CAA and are markedly elevated in moderate and severe CAA (Figure 2. Ba-d). These effects are only observed in the moderate and severe CAA samples and are absent in the mild CAA subgroup, which is not different from controls. The Aβ_40_ peptide is the dominant species whose concentrations in moderate and severe CAA are at least twice and up to 10x increased as compared with other C-terminal forms, respectively. We also noticed the presence of Aβ_41_, albeit in relatively very low quantities, exhibiting similar trends as that of the aforementioned other C-terminal Aβ peptides. Concentrations of Aβ_42_ and Aβ_43_ do not show the same large increase with moderate or severe CAA (Figure 2. Bf-g).

### Contrasting Aβ_1-15_ isomerization (Asp-1, Asp-7) in parenchymal plaques and CAA

We examined the distribution patterns of N-terminal Aβ peptides, specifically isomers of Aβ_1-15_, in parenchymal plaques and CAA. Parenchymal Aβ plaques, vessels without CAA, and vessels with mild CAA showed low levels (∼15%) of native Asp1,7L (native Aβ_1-15_). Moderate and severe CAA groups showed the highest percentage of native Aβ_1-15_ (Figure 2. Ca). While moderate and severe CAA consistently showed a common trend, all comparisons reached statistical significance for the moderate group only. Quantification of the additional isomers showed unique distributions in Aβ plaques, vessels with no or mild CAA, and vessels with moderate or severe CAA. Mono-iso-Asp-L is present at ∼25% across all conditions (Figure 2. Cb). Di-iso-Asp-L is the most common isomer found in Aβ plaques and vessels with no or mild CAA, where it accounts for ∼45% of the Aβ (Figure 2. Cc), while moderate and severe CAA showed lower representation of this isoform (Figure 2. Cd).

The percentage of mono-iso-D-di-iso-Asp found in Aβ plaques (∼15%) and moderate CAA (15%) is significantly lower than in CAA-none and mild groups (∼25%). Although not statistically significant, the percentage of mono-iso-D-di-iso-Asp in severe CAA indicates lower amounts than CAA none and mild CAA.

### Distribution of Aβ N-terminus isoforms in insoluble fractions of (a) CAA and (b) parenchymal Aβ plaques

To ensure all major N-terminal isoforms were accounted for and to investigate the contribution of each isoform in CAA pathology (by severity) and in parenchymal plaque, we calculated the fractional abundance of each isoform over the sum of all isoforms measured. In concordance with the absolute quantitation results above, the fraction of N-terminal Aβ species in CAA is dominated by the presence of Aβ1-15 in mild (94.6%), moderate (98.2%), and severe (97.3%) conditions (Figure 3. Aa). The fractional abundance of N-terminal species in mild CAA is in the order of Aβ_1-15_ >> Aβ_4-15_ > Aβ_3pGlu_ > Aβ_cit-3pGlu_ > Aβ_2-15_. (Refer to SI Table 1 for percentage fractional abundance of Aβ isoforms). However, in the case of moderate CAA, they are in the order of Aβ_1-15_ >> Aβ_3pGlu-15_ > Aβ_4-15_ > Aβ_2-15_ > Aβ_cit-3pGlu-15_ and for severe CAA, Aβ_1-15_ >> Aβ_4-15_ > Aβ_3pGlu-15_ > Aβ_2-15_ > Aβ_cit-3pGlu-15_; In the case of parenchymal plaques, around 48% of the N-terminal species consists of Aβ_1-15_, while Aβ_3pGlu-15_ (∼24%) is the second most abundant species, followed by Aβ_2-15_ (∼14%) and Aβ_4-15_ (∼13%) (Figure 3. Ab). The abundance of Aβ_3pGlu-15_ in our results aligns with the previous results(14) wherein 3pglu was found to be one of the predominant species. Interestingly, the fractional abundance of N-terminal species in parenchymal plaques exhibits far greater heterogeneity than CAA and is in the order of Aβ_1-15_ > Aβ_3pGlu_ > Aβ_2-15_ > Aβ_4-15_.

**Figure 3.**
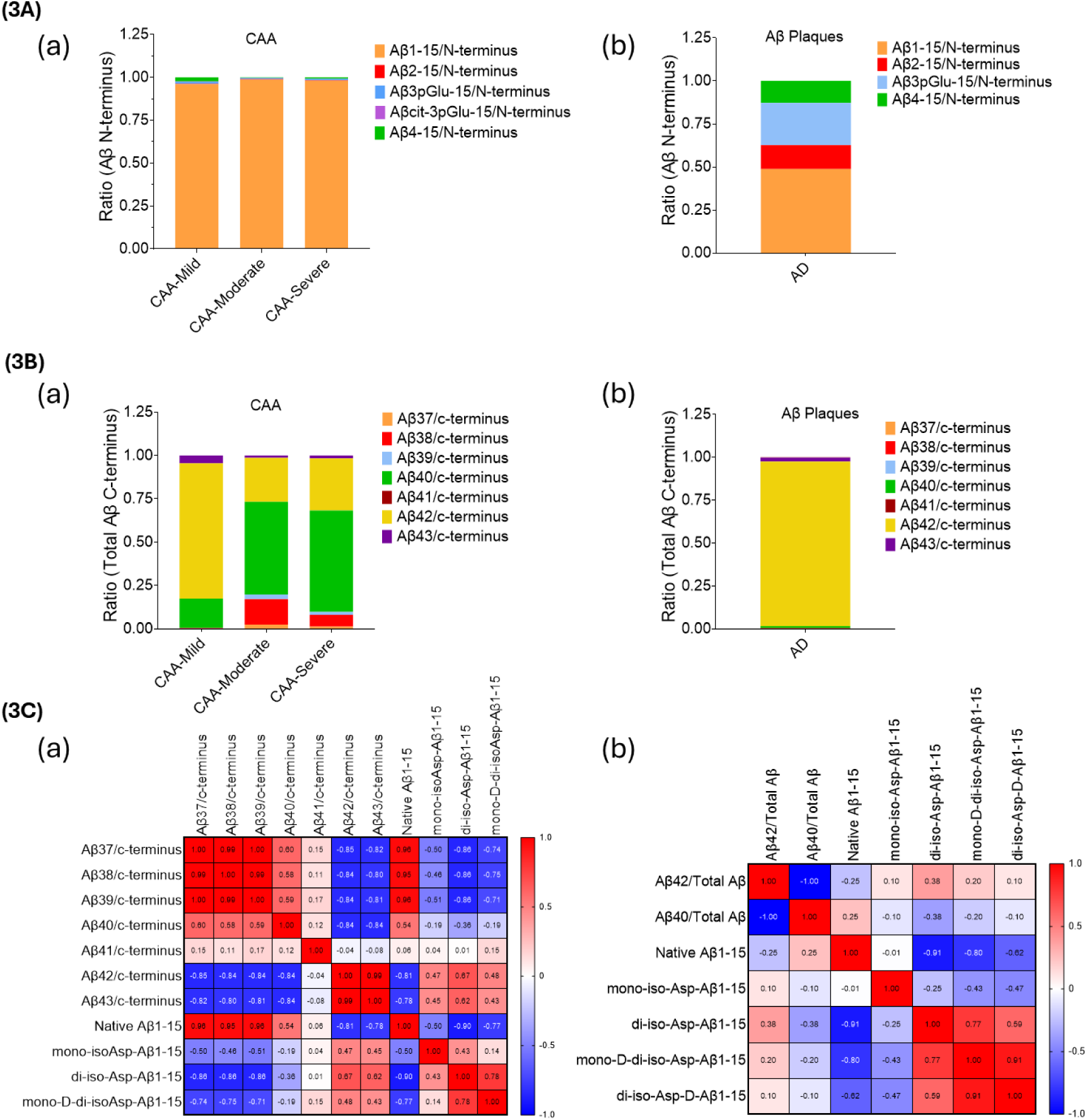
Aβ isoforms of CAA and parenchymal Aβ plaques in insoluble fraction Box plots represent the contribution of each fraction of (3A) N-terminal Aβ isoform in CAA and Aβ plaques. Specifically, (3Aa) in mild, moderate, and severe cases, Aβ1-15 shares the major proportion of 96-99%, while in the case of (3Ab) Aβ parenchymal plaques, Aβ1-15 shares 50% of the proportion. In the case of (3B) C-terminal Aβ isoform in mild CAA consists of ∼78% of Aβ42 in contrast to ∼25% and ∼30% in moderate and severe, respectively. However, the proportion of Aβ40 in mild CAA is ∼17%, while that of moderate and severe CAA is ∼53% and ∼58%. (3C) Comparison of the correlations between Aβ isoforms of (3Ca) CAA and (3Cb) parenchymal Aβ plaques.

**Table 1.** Demographics.

| CAA Severity | Age (years) | Education (years) | ABC score | ApoE Genotype | Sex |
| --- | --- | --- | --- | --- | --- |
| none | 79 | 18 | A0B1C0 | 23 | male |
| none | 101 | 12 | A1B1C0 | 23 | female |
| none | 72 | 16 | A1B0C0 | 33 | female |
| none | 80 | 14 | A1B1C0 | 33 | female |
| none | 72 | 15 | A2B2C0 | 34 | male |
| none | 77 | 20 | A3B3C3 | 34 | male |
| none | 84 | 10 | A3B3C3 | 33 | female |
| none | 102 | 14 | A3B2C1 | 33 | male |
| none | 89 | 12 | A3B2C1 | 34 | female |
| none | 86 | 14 | A3B2C2 | 33 | female |
| mild | 84 | 8 | A3B3C3 | 34 | female |
| mild | 94 | 13 | A3B3C1 | 33 | female |
| mild | 94 | 16 | A3B3C3 | 33 | male |
| mild | 87 | 13 | A3B3C2 | 34 | male |
| mild | 73 | 16 | A3B3C3 | N/A | female |
| moderate | 71 | 16 | A3B3C3 | 44 | female |
| moderate | 83 | 18 | A3B2C1 | 44 | female |
| moderate | 86 | 16 | A3B3C2 | 34 | male |
| moderate | 77 | 15 | A3B3C3 | 44 | female |
| moderate | 85 | 9 | A3B3C3 | 34 | male |
| severe | 93 | 20 | A3B3C3 | 34 | male |
| severe | 93 | 12 | A3B3C2 | 33 | female |
| severe | 93 | 14 | A3B3C3 | 33 | female |
| severe | 78 | 20 | A3B3C3 | 24 | male |
| severe | 88 | 14 | A3B3C3 | 44 | female |

### Distribution of Aβ C-terminus isoforms in insoluble fractions of (a) CAA and (b) parenchymal Aβ plaques

In the case of fractional C-terminal Aβ species, we detected a larger proportion of Aβ_42_ (∼78%) in mild CAA compared to moderate and severe CAA (∼25% and ∼30%, respectively) (Figure S1). In contrast, the proportion of Aβ_40_ in mild CAA was ∼17%, while that in moderate and severe CAA was ∼53% and ∼58%, respectively (Figure 3. Ba) (Proportion of C-terminal Aβ species obtained from the insoluble fractions of individual patient samples has been shown in Figure S3. 3a). With increasing severity of CAA, the fractional contribution of the C-terminus shifted from Aβ_42_ to Aβ_40_. The abundance of fractional C-terminal Aβ species in mild CAA is in the order of Aβ_42_ > Aβ_40_ > Aβ_43_ > Aβ_38_ > Aβ_39_ > Aβ_37_ > Aβ_41_; in moderate CAA it is Aβ_40_ > Aβ_42_ > Aβ_38_ > Aβ_39_ > Aβ_37_ > Aβ_43_ > Aβ_41_; and in severe CAA it is Aβ_40_ > Aβ_42_ > Aβ_38_ > Aβ_43_ > Aβ_39_ > Aβ_37_ > Aβ_41_. In parenchymal plaques, the fractional abundance of C-terminal Aβ species was overwhelmingly dominated by Aβ_42_ (∼96%). The remaining ∼4% consisted of other C-terminal species in the following decreasing order: Aβ_43_ > Aβ_40_ > Aβ_38_ > Aβ_41_ > Aβ_37_ > Aβ_39_ (Figure 3. Bb). Our results indicated high C-terminal heterogeneity in moderate/severe CAA with truncated, less hydrophobic Aβ species (Aβ_37_-Aβ_40_) while Aβ_42_ is the predominant C-terminus in parenchymal plaques.

### Correlation of Aβ C-terminus and N-terminus in CAA and parenchymal Aβ plaques

In samples with CAA, we found a strong positive correlation among shorter C-terminal Aβ species Aβ_37_, Aβ_38_, Aβ_39_, and Aβ_40_ (r=0.58 to 1.00, p value – 0.021 to 2.29 x10-11) (Figure 3. Ca). Additionally, the longer C-terminal Aβ species Aβ_42_ and Aβ_43_ show a strong positive correlation within themselves (r = 0.99, p value < 0.0001). The presence of the shorter C-terminal species (Aβ_37_, Aβ_38_, Aβ_39_, and Aβ_40_) show a strong negative correlation (r = -0.80 to 0.85, p value – 0.0002 to 0.0006) to Aβ_42_ and Aβ_43_ peptides. Additionally, our correlational analysis suggests that C-terminal peptides, Aβ_37_, Aβ_38_, Aβ_39_, and Aβ_40_, exhibited significant positive correlation with native N-terminus Asp1,7L peptide (native Aβ_1-15_) (r = 0.54 - 0.96) The effect was much more pronounced for Aβ_37_ (r = 0.96, p value – 1.24 x 10-7), Aβ_38_ (r = 0.95, p value – 2.8 x 10-7), and Aβ_39_ (r = 0.96, p value – 1.9 x 10-7) than Aβ_40_ (r = 0.54, p value – 0.04). A strong negative correlation between the isomerized N-terminal isoforms (di-iso-Asp-L-Aβ_1-15_ and mono-iso-D-di-iso-Asp-Aβ_1-15_) and the shorter C-terminal isoforms Aβ_37_, Aβ_38_ and Aβ_39_ (r = -0.71 to -0.86, p value – 0.004 to 0.0001) was observed. In parenchymal Aβ plaques, we observed a strong inverse correlation between Aβ_40_ and Aβ_42_ species (r = -1.00, p value – 6 x 10-7) (Figure 3. Cb). We also observed a significant negative correlation (r = -0.77 to -0.90, p value – 0.001 to 1x 10-5) between the native N-terminus Asp1,7L (native Aβ_1-15_) and the N-terminal Asp isomers (di-iso-Asp-Aβ_1-15_ and mono-D-di-iso-Asp-Aβ_1-15_). The N-terminal isomer species (di-iso-Asp-Aβ_1-15_, mono-D-di-iso-Asp-Aβ_1-15_ and di-iso-Asp-D-Aβ_1-15_) show positive correlation with one another (r = 0.59 to 0.91). CAA has lower isomerization and correlates with C-truncated peptides, whereas the plaques with Aβ_42_ have higher isomerized N-terminus.

## Discussion

By employing a vessel purification strategy, coupled with Lys-N digested Aβ-targeted immunoprecipitation/mass spectrometry, we were able to identify the distinct Aβ proteoform signature of CAA, how it changes with disease severity, and how it compares to that of Aβ in parenchymal plaques of AD. Previous studies have demonstrated that Aβ_40_ is the dominant isoform in CAA and Aβ_42_ is the dominant isoform in parenchymal amyloid plaques of AD(20, 24, 34–36). This report is in line with the previously reported Aβ patterns and expands current understanding by providing the first quantitative characterization, including C-terminal isoforms, N-terminal isoforms, and associated PTMs.

One of the main findings of our study was that vast majority of vascular insoluble Aβ exists without any N-terminal truncation in moderate and/or severe CAA. This is in contrast with parenchymal plaque Aβ, where the intact N-terminus represents only 50% of the peptide, with Aβ_2-x_, Aβ_4-x_, and Aβ_3pGlu_ being the other main species (Figure 3. Ab). We additionally measured Aβ Asp1 and Asp7 isomerization in CAA as compared to parenchymal Aβ plaques. CAA aggregates show reduced spontaneous isomerization of Asp1 and Asp7. Although, the precise mechanism for this differential processing remains to be elucidated, such differences in the patterns of PTMs can be explained through the comparison with recently described atomic structures by cryo-electron microscopy (EM) techniques on ex vivo Aβ protofibrils extracted from AD patients.

The structural differences of the protofibril folds of Aβ in CAA vs parenchymal plaques by cryo-EM suggest that modifications of the N-terminus might be access dependent. In parenchymal plaques, the peptides fold in an S-shape such that the N-terminus remains unstructured as part of the fuzzy coat; the disordered N-terminus extends from Asp1-Gly9(37). In contrast, the vascular peptide fold is C-shaped, with its N- and C-termini forming arches and the peptide chain folding back onto the central peptide domain(38). Due to the involvement of the N-terminus in the peptide fold, it follows that solvent accessibility to the N-terminus would be limited in the vascular compartment (less PTMs) as compared to parenchymal plaques.

While our data recapitulated prior work showing that Aβ_40_ is the dominant C-terminal isoform in CAA, we have also measured shorter, less hydrophobic Aβ_37_, Aβ_38_, Aβ_39_ and rarer C-terminal fragments, like Aβ_41_ (although detected in quantities less than 0.1% of total Aβ), the pathological significance of which remains to be elucidated. In characterizing the distribution of C-terminal fragments, we noted a progressive decrease in the fractional abundance of longer, more hydrophobic C-terminal species (Aβ_42_ and Aβ_43_) and an increase in the fractional abundance of shorter, less hydrophobic C-terminal species (Aβ_37_, Aβ_38_, Aβ_39_, and Aβ_40_) with increasing CAA severity. This suggests that hydrophobic C-terminal isoforms, in particular Aβ_42_, might be more functionally relevant to the aggregation process early in the disease course, with Aβ_40_ and other shorter C-terminal fragments becoming dominant later with disease progression and severity in the vasculature. This shift could be compatible with the hypothesis that Aβ_42_ serves as a nidus for the aggregation of Aβ_40_ in the vessels(39). These results highlight that C-terminal heterogeneity is another key distinguishing feature for CAA compared to parenchymal plaques(40). How this proteoform diversity in the vasculature and parenchyma impacts anti-amyloid therapies remains largely unexplored. Critically, it is still unclear which vascular Aβ proteoforms are recognized and targeted by the currently approved anti-Aβ monoclonal antibodies. Adverse side effects such as ARIA continue to pose major challenges to safe and effective treatment for AD. Our analytical platform offers a valuable tool for elucidating the molecular mechanisms underlying ARIA, including the potential involvement of specific vascular Aβ species, and may enable the development of less-invasive biomarkers for stratifying patients at higher risk.

Our combined IP/MS technique expands and complements information provided by structural methods (Cryo-EM) and traditional immunohistochemical studies to further delineate the mechanism by which the same peptide can form distinct fibrils in these two brain compartments. By using mass spectrometry to characterize the proteoform distributions, we have circumvented many of the limitations of immunohistochemistry alone, such as the need for antibodies specific to each truncation or modification. Generally, proteomic investigations lose structural information regarding the large, folded proteins following enzymatic digestion. Our strategy of probing the structural PTMs (Asp residue isomerization) provides new insights into Aβ protofibril folds in distinct brain compartments. Disordered “fuzzy” coat of the aggregates are not always resolved for many disease-associated proteins(41–43). We largely observed a binary response with none and mild-CAA showing similar distributions of Aβ proteoforms and moderate and severe-CAA showing similar distributions. Interestingly, most of the statistically significant relationships were present in only the moderate CAA cohort and did not reach significance for the severe cohort. We believe this is most likely secondary to the variability in CAA pathology within brain regions (occipital, in these experiments) such that tissue submitted for IP/MS analysis may have incompletely represented what was present on routine IHC-stained sections used for grading; and grading itself is a simplified summation of multiple factors (density, vessel burden, and type of vessels affected)(44).

While this report provides unique insights into vascular and parenchymal plaque Aβ, we acknowledge there are limitations to our study. The employed vessel purification protocol excludes larger vessels such as leptomeningeal arteries. We probed CAA from occipital cortex only. Future work using selective capture of vessels stratified by location (leptomeningeal vs. parenchymal), size (arterioles vs. capillaries), or morphology (non-circumferential vs. circumferential vs. dyshoric) will better characterize vessel type-specific proteoform distributions across multiple brain regions. Future studies could also investigate differential changes of N- and C-terminus of intact Aβ in leptomeningeal vs. parenchymal vessels using mass spectrometry techniques such as MALDI-MS imaging(34). Additionally, expanding the proteomic characterization from just the Aβ proteoform distribution to an untargeted proteomic approach could offer additional insights into relevant protein interactions and underlying biological processes in sporadic and familial CAA.

In summary, we provide the first systematic description of the Aβ proteoform distributions across varying degrees of CAA severity, comparing them directly to those found in parenchymal plaques. We show that vascular Aβ in CAA exhibits N-terminal uniformity with minimal Asp isomerization, in contrast to the extensive N-terminal modifications observed in parenchymal plaques, including Asp isomerization, pyroglutamate formation and sequential truncations. Additionally, we identify C-terminal heterogeneity as a defining feature of CAA, marked by enrichment in shorter, less hydrophobic Aβ isoforms (Aβ_37_-Aβ_40_), whereas Aβ_42_ predominates in parenchymal aggregates. We propose that these differences arise from the structural variations in the protofibril folds of Aβ, which influence accessibility and susceptibility of specific regions of peptide to PTMs (Figure 4). Our findings offer insight into the compartment specific Aβ aggregation pathway that could be useful in designing targeted therapeutics aimed at these distinct Aβ species. Moreover, reports indicate lower levels of plasma and CSF Aβ_40_ in CAA diagnosed individuals, implying deposition of Aβ isoforms in the brain vasculature. Detecting these isoforms in the biofluids, particularly the shorter C-terminal species could serve as a valuable biomarker strategy for staging CAA progression and enhancing early diagnosis. Together, our results advance the molecular understanding of Aβ aggregation in CAA and AD and highlight new avenues for both diagnostic and therapeutic development.

**Figure 4.**
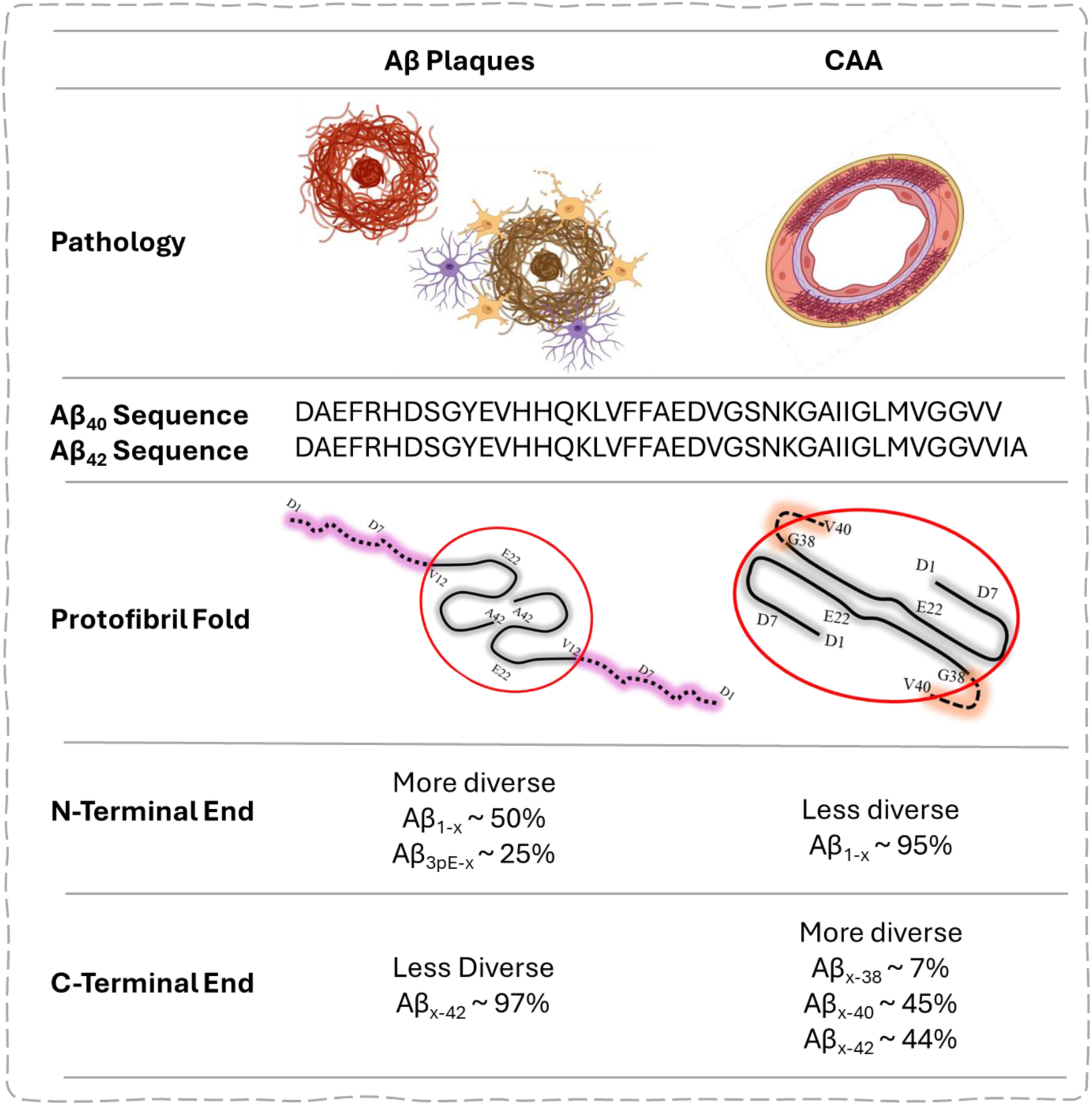
Differences of the protofibril folds of Aβ in CAA vs parenchymal plaques. Dense cored plaque, diffuse plaques, and neuritic plaques in the parenchyma primarily contain protofilaments (red) consisting of G9-A42 or V12-A42 folds (grey ribbon) with the N-terminus being highly disordered (pink dashed region) and prone to PTMs such as Asp isomerization of Asp1/Asp7, while vascular protofibrils (red) contain A1-G38 folds with some disorder in the C-terminus and an ordered N-terminus. Unmodified (native) N-terminus is a key feature of the vascular amyloid in CAA. We hypothesize that Aβ isoform and PTM heterogeneity are thus a consequence of the structure of the protofilaments that are formed in these two distinct aggregates.

## Materials and Methods

### Samples

Autopsy brain samples from AD patients with mild, moderate, or severe CAA (n=5 each), AD patients without CAA (n=5), and age-matched cognitively healthy controls (n=5) were obtained from the Charles F. and Joanne Knight Alzheimer Disease Research Center (Knight ADRC) at Washington University in St. Louis. AD patients without CAA and age-matched cognitively healthy controls were grouped together as the CAA-none cohort. Refer to table 1 for demographics. All participants gave prospective premortem written consent for their brains to be banked and used for research. Data from previously published work regarding Asp1 and Asp7 isomerization of Aβ_1-15_ in parenchymal plaques[27] were used in our analyses.

### Neuropathological assessment

Formalin-fixed autopsy tissue from the left hemisphere of the brain was processed for histochemical and immunohistochemical evaluation. For evaluation of CAA pathology, tissue was stained with antibody 10D5 (Eli Lilly), and sections of occipital cortex were semi quantitatively scored by an expert neuropathologist (none, mild, moderate, and severe) for both leptomeningeal and parenchymal vessels, considering multiple factors including density, circumferential accumulation, and dyshoric features. Sections from the right frozen hemibrain were stained with Methoxy34 immunofluorescence to evaluate side-to-side concordance. Representative images of each CAA category are provided in Figure 1.

### Isolation of cerebral blood vessels and soluble and insoluble fractionation

Cerebral vessels from the postmortem brain samples of AD patients and controls were isolated as described in previous studies[40]. Briefly, 2 g of frozen occipital cortex tissue was homogenized in ice-cold buffer solution at pH 7.4. Next, we added an equal volume of 26% dextran (31390-25G Sigma-Aldrich) to the homogenate and centrifuged at 5800xg for 15 min at 4°C, which separated the samples into three layers: vessel-free parenchyma (upper), myelin/debris (middle), and pellet containing vessels (bottom). The upper and middle layers were discarded. Cerebral vessels from the bottom layer were washed three times with ice-cold buffered solution and further purified by passing over a 100 µm nylon mesh strainer. The filtrate was then centrifuged at 5000×g for 10 min at 4°C to collect cerebrovessels in pellet form.

Isolated cerebral vessels were then suspended in ice-cold 1x RIPA lysis buffer, sonicated, and centrifuged at 20,000xg for 20 min at 4°C, and supernatant was collected as the soluble fraction and stored at –80°C for further experiments. The insoluble amyloid-β fraction was prepared using formic acid as described previously[20, 28]. Briefly, pellets obtained after extraction with RIPA buffer were washed once with PBS, then resuspended in ice-cold 70% formic acid for extraction of insoluble Aβ. The formic acid fraction was neutralized in 20 volumes of 1M Tris solution. The protein concentration in the supernatant from the insoluble fraction was determined using the DC protein assay kit from Bio-Rad (Cat # 5000001).

### Immunoprecipitation/Mass Spectrometry

For the soluble and insoluble Aβ peptide analysis, we used the HJ5.1 antibody for immunoprecipitation (IP) of Aβ. Each brain sample was normalized to a total tissue weight of 10 µg and diluted to 500 µL using 0.01% human serum albumin (has) in milli-Q water before adding antibody coupled beads. 20 µL of recombinant 15NAβ internal standard (0.1 ng/µL Aβ_1-42_, 0.01 ng/µL Aβ_1-40_, 0.01 ng/µL Aβ_1-43_, and 0.01 ng/µL Aβ_1-38_ prepared in 0.1% NH_4_OH/20% ACN) was spiked into the samples. 50 µL of HJ5.1-coupled DynaBeads (25 µg antibody/mg beads per IP) and 25 µL of 1% IGEPAL (I8896-100ML, Sigma-Aldrich), 1x protease inhibitor, 5 mM guanidine in PBS pH 7.4 were added to the samples. The IP and bead washing steps were performed in a KingFisher automated format; Aβ peptides were eluted off beads using neat formic acid (99% FA) and lyophilized to dryness. The samples were resuspended in 80 µL of 100 mM TEABC, and 20 µL of Lys-N enzyme (2.5 ng/µL) was added for overnight digestion at 37℃. The following day the peptides were desalted using an Oasis HLB µElution plate (Waters) according to the manufacturers’ protocol and lyophilized to dryness. During the desalting process, 10 fmol each of AQUA internal-standard (Thermo Fisher Scientific) peptide for Aβ_1-15_, Aβ_2-15_, Aβ_3pGlu-15_, and Aβ_4-15_ were spiked for quantification. The peptides were incubated in 3% H₂O₂ and 3% FA for 18 hours at 4℃ to oxidize the methionine within Aβ peptides. The following day, samples were desalted using an Oasis µElution HLB plate (Waters) using the manufacturers’ protocol and lyophilized. The Aβ peptides were reconstituted in a 2%ACN/2%FA solution containing 20 fmol/µL BSA tryptic peptides for MS analysis on a Vanquish Neo UHPLC (Thermo Fisher Scientific) coupled to Orbitrap Exploris 480 (Thermo Fisher Scientific).

The peptides were directly loaded onto HSS T3 75 μm × 100 μm, 1.8 μm C18 column (Waters (heated to 65°C)) using a Vanquish Neo UHPLC (Thermo Fisher Scientific, San Jose, CA, USA) with a flow rate of 0.7 µL/min using buffer A (0.1 % FA in water) and buffer B (0.1% FA in acetonitrile). The Aβ peptides were eluted off the column using a flow rate of 0.4 µL/min with a gradient of buffer B of 0.5-4% for 2 min, 4-8% for 5 min, 8-20% for 10 min, and 20-35% for 3 minutes before ramping to 90% buffer B and cleaning the column for the next 3 min. The column was equilibrated back to 0.5% buffer B for the next 4 min before starting another run. The Orbitrap Exploris 480 was equipped with a Nanospray Flex electrospray ion source (Thermo Fisher Scientific, San Jose, CA, USA) and operated in positive ion mode. Peptide ions were sprayed from a 10 μm SilicaTip emitter (New Objective, Woburn, MA, USA) into the ion source (spray voltage = 2200 V, ion source temperature 275°C). Peptide precursor ions were targeted and isolated in the quadrupole. Isolated ions were fragmented by HCD, and fragment ions were detected in the Orbitrap (resolution of 30,000–60,000 for Aβ peptides, mass range 150–1,500 m/z). The raw MS data were imported into Skyline (University of Washington, v4.2) with formula annotations of the targeted peptides. Aβ peptides (Aβ_1-15_, Aβ_2-15_, Aβ_4-15_, Aβ_3pGlu-15_, Aβ_16-27_, Aβ_28-37_, Aβ_28-38_, Aβ_28-39_, Aβ_28-40_, Aβ_28-41_, Aβ_28-42_, Aβ_28-43_) were quantified by comparison with the corresponding isotopomer signals from their respective 15N internal standard (15N Aβ_1-15_ peak area was used to quantitate Aβ_2-15_, Aβ_4-15_, Aβ_3pGlu-15_, 15N Aβ_28-38_, 15N Aβ_28-40_, and 15N Aβ_28-42_ and were used to quantitate Aβ_28-37_, Aβ_28-39_, and Aβ_28-41_, respectively).

### Statistical analysis

Data analyses were performed in Tableau, GraphPad Prism and R. Statistical analysis of the Aβ isomers, isoforms and their fractional values of each group were calculated using non-parametric Kruskal Wallis test followed by post-hoc pairwise multiple comparison using the two-stage method of Benjamini, Kreiger, and Yekutieli. P values were adjusted for multiple comparison false discovery rate (P<0.05). For correlation heatmaps of Aβ species, non-parametric Spearman correlations were performed (confidence interval of 95%).

### Significance Statement

Aggregation and deposition of amyloid-β (Aβ) is a hallmark of cerebral amyloid angiopathy (CAA) and Alzheimer’s disease (AD). The molecular mechanisms driving the compartment-specific deposition of Aβ in brain vasculature and extracellular matrix, particularly those involving isomerization and post translational modifications, are poorly understood. This study highlights how brain-derived Aβ proteoforms differ between mild, moderate, and severe CAA, and in parenchymal plaques. We hypothesize that these differences may be due to distinct structural folds in Aβ aggregates, influencing susceptibility and accessibility to isomerization and PTMs. Understanding the biochemistry and compartment-specific aggregation patterns supports the development of targeted therapies. Moreover, detecting these isoforms in blood or CSF could aid in the staging and early diagnosis of CAA.

## Acknowledgments

We thank the participants and personnel of the Knight Alzheimer Disease Research Center, as well as the staff of Washington University’s Translational Human Neurodegenerative Disease Research (THuNDR) Laboratory for providing postmortem human brain tissue for this study. We thank Vitaliy Ovod for helping in data analysis on Tableau software.

## Data Availability Statement

The quantitative data is available as supplementary materials accompanying this article. Additional data and materials supporting the findings of the study can be requested from the corresponding authors.

## Ethics and Consent to Participate declarations

Autopsy samples were obtained from the Knight Alzheimer’s Disease Research Center at Washington University in St. Louis. All participants gave prospective premortem written consent for their brains to be banked and used for research.

## Author Contributions

Conceptualization: S.M. and K.E.S.; Investigation: J.M., S.M. and R.A.C.; Formal Analysis: S.M., K.F.R., and S.K.; Writing – original draft preparation: K.F.R. and S.K.; Writing – review and editing: S.M., K.E.S., G.J.Z., C.S., J.M. and R.J.B.; Funding acquisition: S.M. and K.E.S.; Resources: R.J.B., S.M., J.M. and C.S.; Supervision: R.J.B., G.J.Z., J.M., K.E.S. and S.M. All authors have reviewed the manuscript and provided their consent for publication.

## References

1. C. L. Masters, et al., Amyloid plaque core protein in Alzheimer disease and Down syndrome. Proc Natl Acad Sci U S A 82, 4245–4249 (1985).

2. M. Goedert, M. G. Spillantini, R. Jakes, D. Rutherford, R. A. Crowther, Multiple isoforms of human microtubule-associated protein tau: sequences and localization in neurofibrillary tangles of Alzheimer’s disease. Neuron 3, 519–526 (1989).

3. L. Walker, H. Simpson, A. J. Thomas, J. Attems, Prevalence, distribution, and severity of cerebral amyloid angiopathy differ between Lewy body diseases and Alzheimer’s disease. Acta Neuropathol Commun 12, 28 (2024).

4. M. Yamada, Cerebral amyloid angiopathy: an overview. Neuropathology 20, 8–22 (2000).

5. H. A. D. Keage, et al., Population studies of sporadic cerebral amyloid angiopathy and dementia: a systematic review. BMC Neurol 9, 3 (2009).

6. A. E. Roher, et al., Structural alterations in the peptide backbone of beta-amyloid core protein may account for its deposition and stability in Alzheimer’s disease. J Biol Chem 268, 3072–3083 (1993).

7. J. Attems, K. Jellinger, D. R. Thal, W. Van Nostrand, Review: Sporadic cerebral amyloid angiopathy. Neuropathology Appl Neurobio 37, 75–93 (2011).

8. M. A. Mintun, et al., Donanemab in Early Alzheimer’s Disease. N Engl J Med 384, 1691– 1704 (2021).

9. C. H. Van Dyck, et al., Lecanemab in Early Alzheimer’s Disease. N Engl J Med 388, 9–21 (2023).

10. R. Vassar, et al., Beta-secretase cleavage of Alzheimer’s amyloid precursor protein by the transmembrane aspartic protease BACE. Science 286, 735–741 (1999).

11. Department of Psychiatry and Psychotherapy, University Medical Center (UMG), Georg-August-University, Göttingen, Germany, et al., “N-Terminally Truncated Aß Peptide Variants in Alzheimer’s Disease” in Alzheimer’s Disease, New York University Alzheimer’s Disease Center, New York University School of Medicine, New York, USA, T. Wisniewski, Eds. (Codon Publications, 2019), pp. 107–122.

12. W. He, C. J. Barrow, The Aβ 3-Pyroglutamyl and 11-Pyroglutamyl Peptides Found in Senile Plaque Have Greater β-Sheet Forming and Aggregation Propensities in Vitro than Full-Length Aβ. Biochemistry 38, 10871–10877 (1999).

13. E. Cabrera, et al., Aβ truncated species: Implications for brain clearance mechanisms and amyloid plaque deposition. Biochimica et Biophysica Acta (BBA) - Molecular Basis of Disease 1864, 208–225 (2018).

14. A. E. Roher, et al., beta-Amyloid-(1-42) is a major component of cerebrovascular amyloid deposits: implications for the pathology of Alzheimer disease. Proc Natl Acad Sci U S A 90, 10836–10840 (1993).

15. N. Kakuda, et al., Distinct deposition of amyloid-β species in brains with Alzheimer’s disease pathology visualized with MALDI imaging mass spectrometry. Acta Neuropathol Commun 5, 73 (2017).

16. D. L. Miller, et al., Peptide compositions of the cerebrovascular and senile plaque core amyloid deposits of Alzheimer’s disease. Arch Biochem Biophys 301, 41–52 (1993).

17. E. Gkanatsiou, et al., A distinct brain beta amyloid signature in cerebral amyloid angiopathy compared to Alzheimer’s disease. Neuroscience Letters 701, 125–131 (2019).

18. E. Portelius, et al., Mass spectrometric characterization of brain amyloid beta isoform signatures in familial and sporadic Alzheimer’s disease. Acta Neuropathol 120, 185–193 (2010).

19. B. Lyons, M. Friedrich, M. Raftery, R. Truscott, Amyloid Plaque in the Human Brain Can Decompose from Aβ(1-40/1-42) by Spontaneous Nonenzymatic Processes. Anal Chem 88, 2675–2684 (2016).

20. W. Michno, et al., Pyroglutamation of amyloid-βx-42 (Aβx-42) followed by Aβ1–40 deposition underlies plaque polymorphism in progressing Alzheimer’s disease pathology. Journal of Biological Chemistry 294, 6719–6732 (2019).

21. T. C. Saido, et al., Dominant and Differential Deposition of Distinct I -Amyloid Peptide Species, AI∼N3(pE), in Senile Plaques.

22. Y.-M. Kuo, M. R. Emmerling, A. S. Woods, R. J. Cotter, A. E. Roher, Isolation, Chemical Characterization, and Quantitation of Ab 3-Pyroglutamyl Peptide from Neuritic Plaques and Vascular Amyloid Deposits. BIOCHEMICAL AND BIOPHYSICAL RESEARCH COMMUNICATIONS 237 (1997).

23. S. Mukherjee, et al., Citrullination of Amyloid-β Peptides in Alzheimer’s Disease. ACS Chem Neurosci 12, 3719–3732 (2021).

24. S. Mukherjee, et al., Quantification of N-terminal amyloid-β isoforms reveals isomers are the most abundant form of the amyloid-β peptide in sporadic Alzheimer’s disease. Brain Communications 3, fcab028 (2021).

25. Y. Tomidokoro, et al., Iowa variant of familial Alzheimer’s disease: accumulation of posttranslationally modified AbetaD23N in parenchymal and cerebrovascular amyloid deposits. Am J Pathol 176, 1841–1854 (2010).

26. Y. Wakutani, et al., Novel amyloid precursor protein gene missense mutation (D678N) in probable familial Alzheimer’s disease. J Neurol Neurosurg Psychiatry 75, 1039–1042 (2004).

27. Y. Shin, et al., Aβ species, including IsoAsp23 Aβ, in Iowa-type familial cerebral amyloid angiopathy. Acta Neuropathol 105, 252–258 (2003).

28. Y. Tomidokoro, et al., Iowa Variant of Familial Alzheimer’s Disease. The American Journal of Pathology 176, 1841–1854 (2010).

29. S. Fossati, K. Todd, K. Sotolongo, J. Ghiso, A. Rostagno, Differential contribution of isoaspartate post-translational modifications to the fibrillization and toxic properties of amyloid β and the Asn23 Iowa mutation. Biochem J 456, 347–360 (2013).

30. S. Schrempel, et al., Identification of isoAsp7-Aβ as a major Aβ variant in Alzheimer’s disease, dementia with Lewy bodies and vascular dementia. Acta Neuropathol 148, 78 (2024).

31. S. Mukherjee, et al., Isomerized Aβ in the brain can distinguish the status of amyloidosis in the Alzheimer’s Disease spectrum. [Preprint] (2025). Available at: http://biorxiv.org/lookup/doi/10.1101/2025.04.29.650793 [Accessed 08 May 2025].

32. T. R. Lambeth, et al., Spontaneous Isomerization of Long-Lived Proteins Provides a Molecular Mechanism for the Lysosomal Failure Observed in Alzheimer’s Disease. ACS Cent Sci 5, 1387–1395 (2019).

33. H. Crehan, et al., Effector function of anti-pyroglutamate-3 Aβ antibodies affects cognitive benefit, glial activation and amyloid clearance in Alzheimer’s-like mice. Alz Res Therapy 12, 12 (2020).

34. W. Michno, et al., Chemical traits of cerebral amyloid angiopathy in familial British-, Danish-, and NON-ALZHEIMER ’s dementias. Journal of Neurochemistry 163, 233–246 (2022).

35. A. Rostagno, E. Cabrera, T. Lashley, J. Ghiso, N-terminally truncated Aβ4-x proteoforms and their relevance for Alzheimer’s pathophysiology. Transl Neurodegener 11, 30 (2022).

36. S. Koutarapu, et al., Chemical imaging delineates Aβ plaque polymorphism across the Alzheimer’s disease spectrum. Nat Commun 16, 3889 (2025).

37. Y. Yang, et al., Cryo-EM structures of amyloid-β 42 filaments from human brains. Science 375, 167–172 (2022).

38. M. Kollmer, et al., Cryo-EM structure and polymorphism of Aβ amyloid fibrils purified from Alzheimer’s brain tissue. Nat Commun 10, 4760 (2019).

39. R. F. Sowade, T. R. Jahn, Seed-induced acceleration of amyloid-β mediated neurotoxicity in vivo. Nat Commun 8, 512 (2017).

40. J. M. Fulcher, et al., Discovery of Proteoforms Associated With Alzheimer’s Disease Through Quantitative Top-Down Proteomics. Molecular & Cellular Proteomics 24, 100983 (2025).

41. Y. Yang, et al., Cryo-EM structures of Aβ40 filaments from the leptomeninges of individuals with Alzheimer’s disease and cerebral amyloid angiopathy. Acta Neuropathol Commun 11, 191 (2023).

42. S. H. W. Scheres, B. Ryskeldi-Falcon, M. Goedert, Molecular pathology of neurodegenerative diseases by cryo-EM of amyloids. Nature 621, 701–710 (2023).

43. S. Lövestam, et al., Disease-specific tau filaments assemble via polymorphic intermediates. Nature 625, 119–125 (2024).

44. J. Attems, Sporadic cerebral amyloid angiopathy: pathology, clinical implications, and possible pathomechanisms. Acta Neuropathol 110, 345–359 (2005).

