## Supplementary material for "Post-translational modifications distinguish amyloid-β isoform patterns extracted from vascular deposits and parenchymal plaques": BioRxiv SI.docx

**Supplementary Information (SI)**

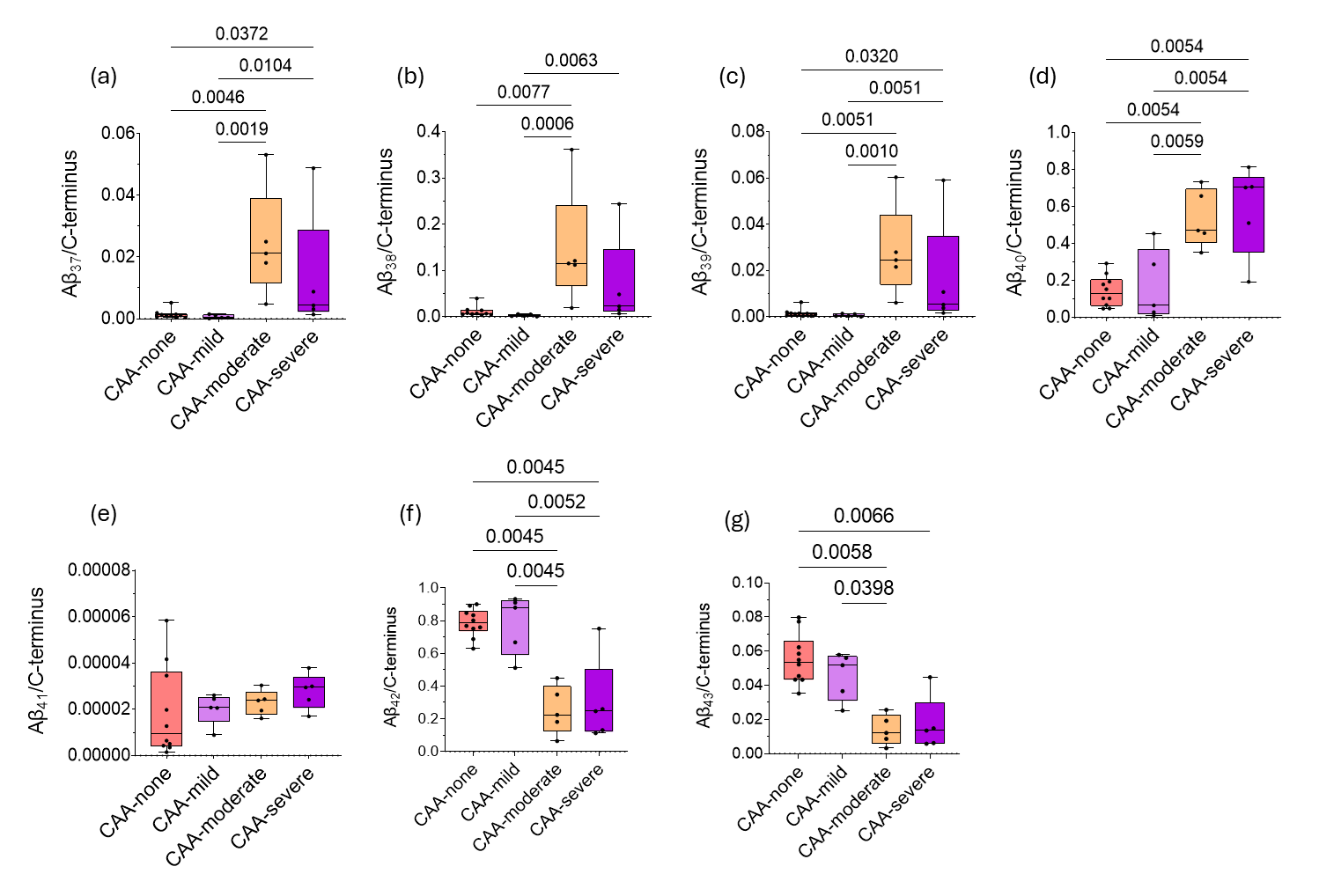

**S1.** Boxplots of fractional C-terminal Aβ species in the insoluble fraction of mild, moderate, and severe CAA.

**Hydrophilic Aβ species show increased proportions, while hydrophobic Aβ species show decreased proportions** **in moderate and severe CAA: fractional Aβ C-terminus contribution in the vessel insoluble preparation.**

The calculation of the fractional contribution of each C-terminal species (Aβ_37_, Aβ_38_, Aβ_39_, Aβ_42_, and Aβ_43_) among all C-terminal truncated Aβ species shows a significant increase, particularly of Aβ_37_, Aβ_38_, and Aβ_39_ proportions in moderate and severe CAA. However, among the hydrophilic species, the fold change of Aβ_38_ and Aβ_40_ species with CAA severity is lower in mild conditions, which profoundly increases in moderate and severe conditions in comparison with Aβ_37_, Aβ_38_, and Aβ_39_ species. Interestingly, we also noticed that the fractions of Aβ_42_ and Aβ_43_ species decrease with CAA severity, an important factor that distinguishes vascular from parenchymal amyloid composition. Higher levels are found in mild conditions, while these proportions significantly drop in moderate and severe conditions.

**Aβ N-terminus in the Soluble Fraction**

We then investigated the differences between N-terminal and C-terminally truncated forms of Aβ species in the brain RIPA buffer soluble fraction (SI Fig 2A). Our analysis of Aβ N-terminus in the soluble fraction revealed the presence of Aβ_1-15_ and Aβ_3pGlu-15_, wherein the concentration of Aβ_1-15_ was found to be 100 times higher than the N-truncated Aβ_3pGlu-15_ species in CAA-positive samples.

**Aβ C-terminus in soluble fraction shows marked elevation in moderate and severe CAA**

Our results show an increase in the concentrations of Aβ_37_, Aβ_38_, Aβ_39_, and Aβ_40_ with the severity of CAA, which are markedly elevated in moderate and severe CAA (SI Fig 2B). Specifically, Aβ_40_ exhibits lower levels in mild CAA, which increases by 3-5 folds in moderate and severe CAA. Like the insoluble fraction, our results indicate higher levels of Aβ_42_ in mild conditions, which drop by over 2 folds in moderate and severe conditions.

**Distribution of Aβ C-terminus isoforms in soluble fractions of CAA**

In the case of the soluble fraction of CAA, we detected a larger proportion of Aβ_42_ (~83%) followed by Aβ_40_ (~14%) in mild CAA (SI Fig 2C). In moderate and severe CAA, we noticed equivalent proportions of Aβ_40_ and Aβ_42_. Specifically, we observed ~45% and ~47% of Aβ_40_ in moderate and severe CAA, respectively. While the fractional abundance of Aβ_42_ in moderate and severe was found to be ~43% and ~46%, respectively. Our results also indicate higher proportions of Aβ_38_ in moderate (~9%) than in severe CAA (~4%).

**Correlation of Aβ C-terminus isoforms and N-terminus in CAA — Whole Cohort-Soluble.**

We found a *significant positive correlation* among C-terminal Aβ species Aβ_37_, Aβ_38_, Aβ_39_, and Aβ_40_ (r value ranging from 0.75 to 0.82, p value 1.37x10^-5^ to 3.4x10^-29^), the correlation between Aβ_42_ and Aβ_43_ peptides in the soluble fraction is not significant (SI Fig 2D). However, the relation between C-terminus Aβ species Aβ_37_, Aβ_38_, Aβ_39_, Aβ_40_ and Aβ_42_ peptides indicate a significant negative correlation (r = -0.67 to -0.89, p value 2.3x10^-4^ to 4.1x10^-9^), while there is no correlation with Aβ_43_. Additionally, our correlational analysis suggests that C-terminus Aβ_37_, Aβ_38_, Aβ_39_, and Aβ_40_ peptides exhibit significant *positive correlation* with the (Native Aβ_1-15_) (r = 0.42 to 0.58, p Value 3.7x10^-2^ to 5.4x10^-3^) and (mono-D-di-isoAsp-Aβ_1-15_) (r = 0.53 to 0.73), while the (mono-isoAsp-Aβ_1-15_) indicates negative correlation (r = -0.40 to -0.63). However, Aβ_42_ exhibits a positive correlation with (mono-isoAsp-Aβ_1-15_ (r = 0.47, p value 1.8x10^-2^) and negative correlation with mono-D-di-isoAsp-Aβ_1-15_ (r = - 0.49, p value 1,5x10^-2^). Aβ_43_ indicates positive correlation with di-iso-Asp-Aβ_1-15_ (r = 0.40, p value 0.047)_._

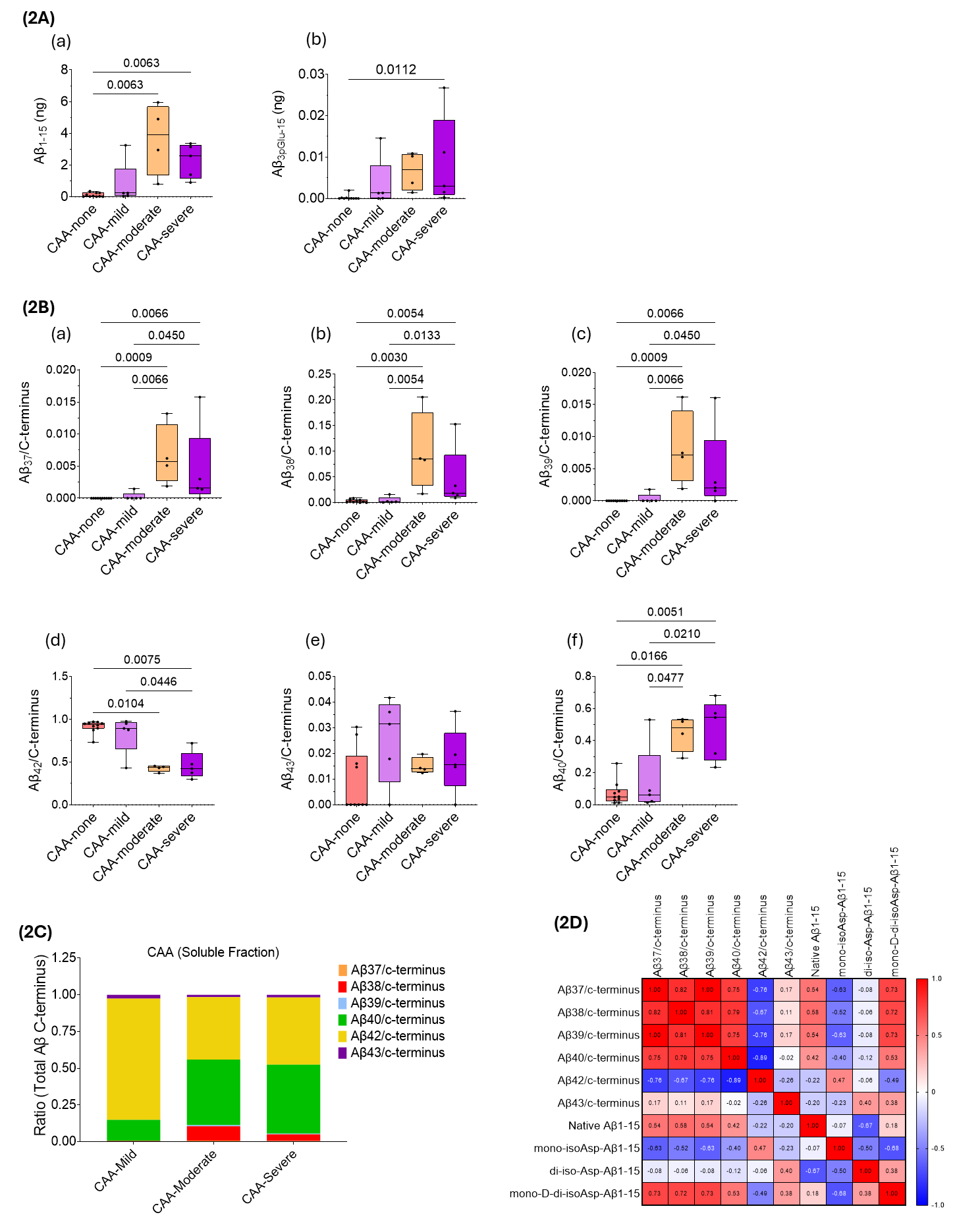

**S2.** **Soluble fraction of mild, moderate, and severe CAA**

Box plots of the concentration of (2A) N-terminal and (2B) fractional C- terminal Aβ species in the soluble fraction of mild, moderate, and severe CAA. (2C) Proportion of C-terminal Aβ species obtained from the soluble fractions of mild, moderate, and severe CAA and (2D) Spearman correlation plot of the Aβ isoforms.

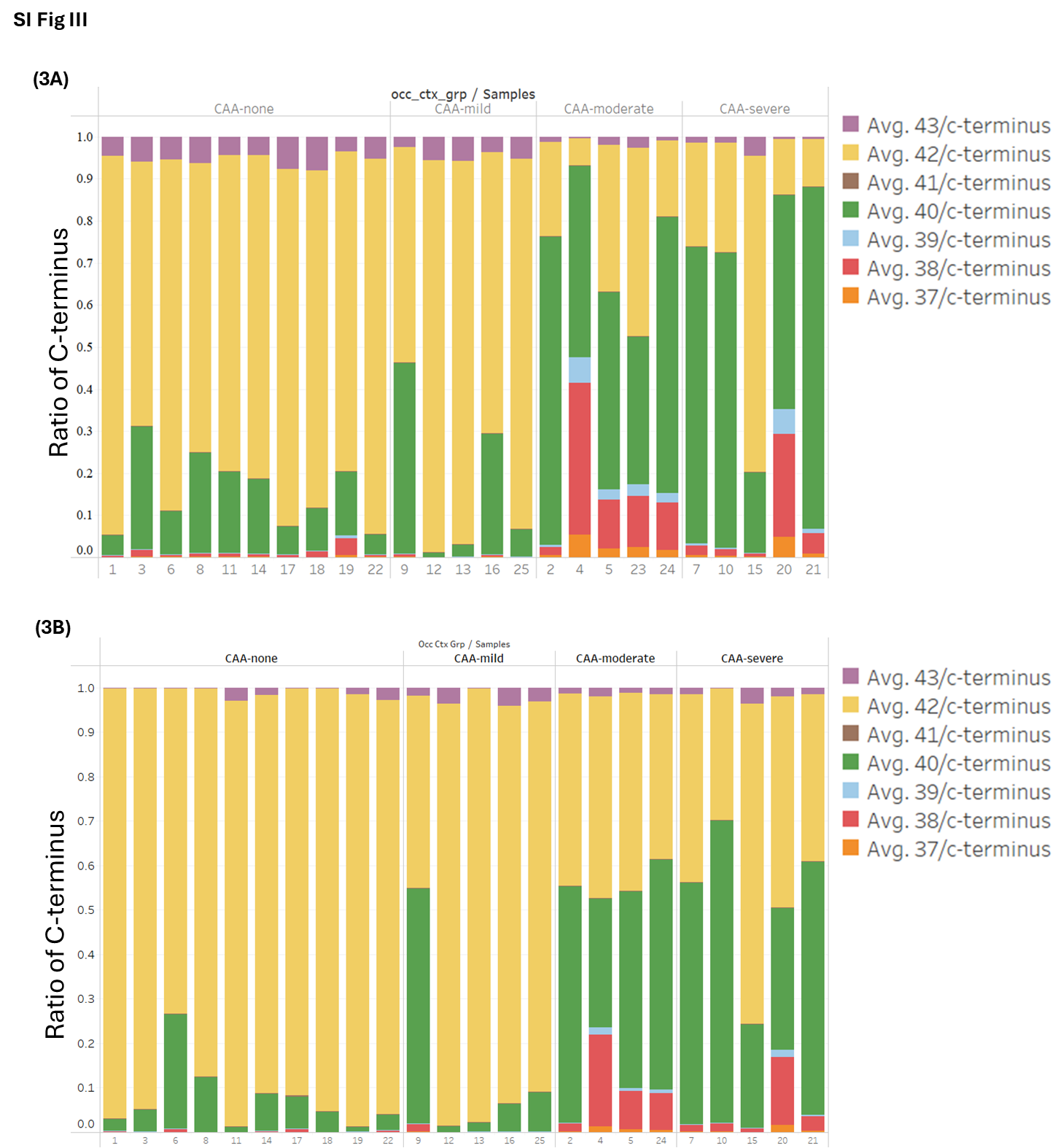

**S3. Insoluble and soluble fractions of individual patient samples**

Proportion of C-terminal Aβ species obtained from the (3A) insoluble and (3B) soluble fractions of individual patient samples (N=25).

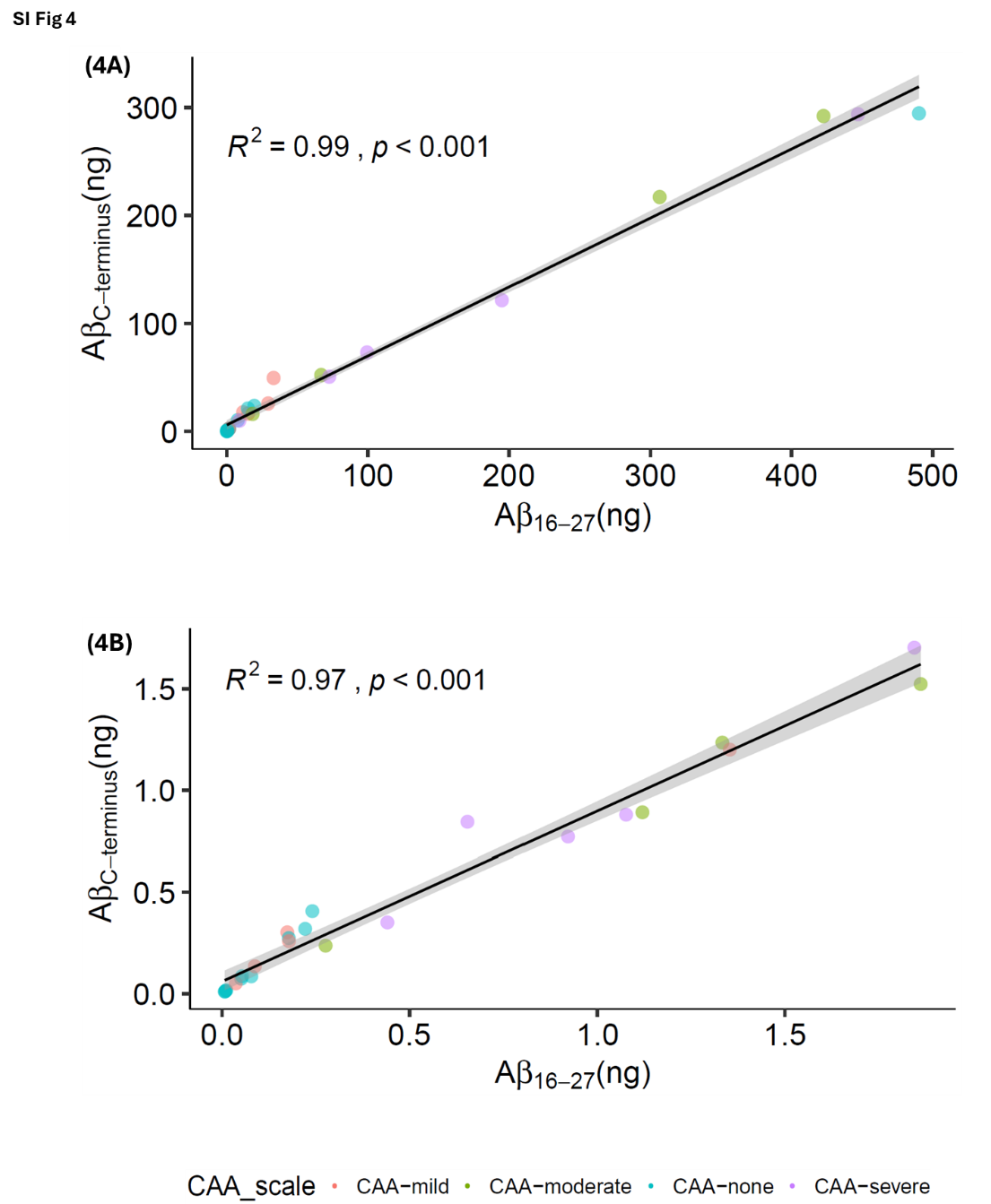

**S4. Insoluble (4A) and soluble (4B) C-terminus comparison with mid-domain Aβ.** Comparison of the sum of the concentration of C-terminal species with the concentration (ng) of the mid-domain Aβ species (Aβ_16-27_). We observed a strong correlation in the concentration of total C-terminal Aβ species and the concentration of mid-domain Aβ both in the insoluble (R^2^ = 99, p < 0.001) and soluble fraction (R^2^ = 0.97, p < 0.001). This highlights the sum of all the major C-terminal peptides measured after mid-domain IP (HJ5.1) captures total Aβ present in these samples.

**SI Table 1: Order of fractional abundance of Aβ species in CAA and parenchymal plaques**

|  | **The fractional abundance of Aβ species** | |
| --- | --- | --- |
| **N-terminal Aβ species in insoluble fractions** | | |
| Mild CAA | | Aβ_1-15_ (~96%) >> Aβ_4-15_ (~2.3%) > Aβ_3pGlu-15_ (~1.2%)  >Aβ_cit-3pGlu-15_ (~0.3%) > Aβ_2-15_(~0.2%) |
| Moderate CAA | | Aβ_1-15_ (~99%) >> Aβ_3pGlu-15_ (~0.6%) > Aβ_4-15_ (~0.3%)  > Aβ_2-15_(~0.2%) > Aβ_cit-3pGlu-15_ (~0.1%) |
| Severe CAA | | Aβ_1-15_ (~98%) >> Aβ_4-15_ (~0.8%) > Aβ_3pGlu-15_ (~0.8%)  > Aβ_2-15_(~0.2%) > Aβ_cit-3pGlu-15_ (~0.05%) |
| Parenchymal Plaques | | Aβ_1-15_ (~49%) > Aβ_3pGlu-15_ (~25%) > Aβ_2-15_ (~14%) > Aβ_4-15_(~13%) |
| **C-terminal Aβ species in insoluble fractions** | | |
| Mild CAA | | Aβ_42_ (~78%) >> Aβ_40_ (~17%) > Aβ_43_ (~5%) > Aβ_38_ (~0.2%)  > Aβ_39_ (~0.07%) > Aβ_37_ (~0.06%) > Aβ_41_(~0.002%) |
| Moderate CAA | | Aβ_40_ (~53%) >> Aβ_42_ (~25%) > Aβ_38_ (~15%) > Aβ_39_ (~3%)  > Aβ_37_ (~2.4%) > Aβ_43_ (~1.3%) > Aβ_41_(~0.002%) |
| Severe CAA | | Aβ_40_ (~59%) >> Aβ_42_ (~30%) > Aβ_38_ (~7%) > Aβ_43_ (~1.7%)  > Aβ_39_ (~1.6%) > Aβ_37_ (~1.3%) > Aβ_41_(~0.003%) |
| Parenchymal Plaques | | Aβ_42_ (~96%) >> Aβ_43_ (~2.4%) > Aβ_40_ (~1.0%) > Aβ_38_ (~0.3%)  > Aβ_41_ (~0.1%) > Aβ_37_ (~0.07%) > Aβ_39_ (~0.009%) |
| **C-terminal Aβ species in soluble fractions** | | |
| Mild CAA | | Aβ_42_ (~83%) >> Aβ_40_ (~14.3%) > Aβ_43_ (~2.5%) > Aβ_38_ (~0.4%)  > Aβ_39_ (~0.03%) > Aβ_37_ (~0.03%) |
| Moderate CAA | | Aβ_40_ (~45%) > Aβ_42_ (~43%) > Aβ_38_ (~10%) > Aβ_43_ (~1.5%)  > Aβ_39_ (~0.8%) > Aβ_37_ (~0.7%) |
| Severe CAA | | Aβ_40_ (~47%) > Aβ_42_ (~46%) > Aβ_38_ (~4.6%) > Aβ_43_ (~1.7%)  > Aβ_39_ (~0.4%) > Aβ_37_ (~0.4%) |
